# Massively parallel clonal analysis using CRISPR/Cas9 induced genetic scars

**DOI:** 10.1101/056499

**Authors:** Jan Philipp Junker, Bastiaan Spanjaard, Josi Peterson-Maduro, Anna Alemany, Bo Hu, Maria Florescu, Alexander van Oudenaarden

## Abstract

A key goal of developmental biology is to understand how a single cell transforms into a full-grown organism consisting of many cells. Although impressive progress has been made in lineage tracing using imaging approaches, analysis of vertebrate lineage trees has mostly been limited to relatively small subsets of cells. Here we present scartrace, a strategy for massively parallel clonal analysis based on Cas9 induced genetic scars in the zebrafish.

---

The timing of each cell division and the fate of the progeny define the lineage of an organism. Analysis of the lineage history of cell populations can reveal the developmental origin and the clonality of cell populations^1^. Genetically encoded fluorescent proteins are widely used as lineage markers^2,3^, but due to limited spectral resolution this approach has mostly been restricted to tracking the lineage of a relatively small subset of cells. Recent progress in live imaging has allowed for following many individual cells over time in optically transparent samples such as early fly and zebrafish embryos^4,5^. Nevertheless, direct observation of all cell divisions is generally only possible at the earliest developmental stages. RNA sequencing has emerged as a powerful method for systematic expression profiling of single cells and for computational inference of differentiation dynamics^6-8^. However, our ability to harness the enormous multiplexing capacity of high-throughput sequencing for lineage tracing has so far been lagging behind, despite pioneering studies using somatic mutations^9^, transposon tagging^10^, or viral barcoding^11,12^. While these techniques have yielded important insights into developmental lineage decisions, they are typically limited to tracing the lineage of a small subset of cells within the organism. Here we present scartrace, a method for massively parallel clonal analysis on the level of the whole organism. In scartrace we use CRISPR/Cas9 genome editing for clonal analysis, in an approach that is related to the recently published GESTALT method^13^.

Scartrace is based on the observation that, in the absence of a template for homologous repair, Cas9 produces short insertions or deletions (indels) at its target sites, which are variable in their length and position^14,15^. We reasoned that these indels (hereafter referred to as genetic “scars”) constitute a permanent, heritable cellular barcode that can be used for lineage analysis. To ensure that genetic scarring does not interfere with the viability of the cells, we targeted GFP in a zebrafish line with a histone-GFP transgene^16^. We injected sgRNA for GFP and Cas9 mRNA or protein into 1-cell stage embryos in order to mark individual cells with genetic scars at an early time point in development (Fig. 1a). Loss of GFP fluorescence in injected embryos served as a direct visual confirmation of efficient scar formation. Scars were then analyzed at a later time by targeted sequencing of GFP (see Materials and Methods).

**Figure 1.**
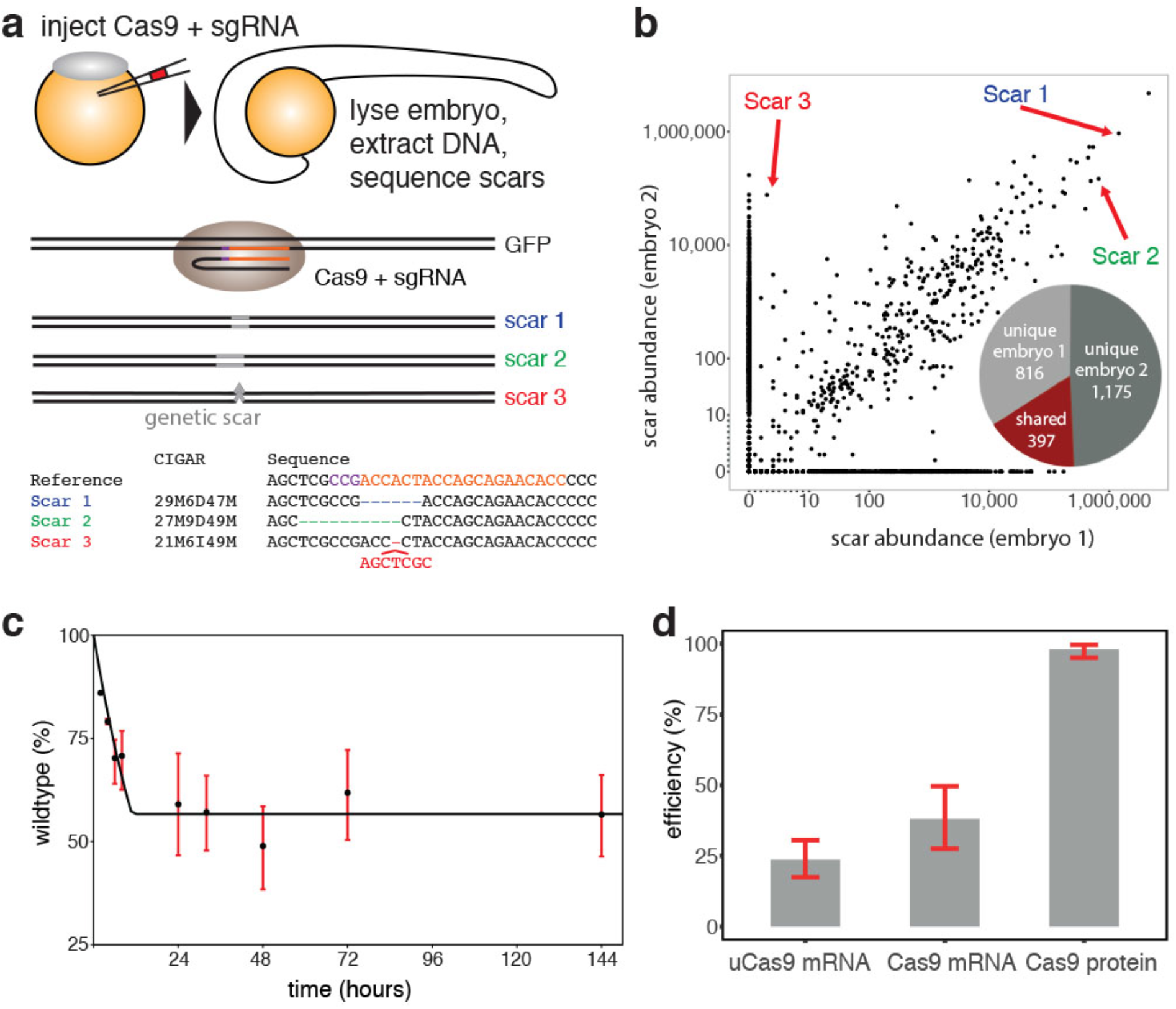
Cas9 generates large diversity of single-cell barcodes. (**a**). Sketch of the experimental protocol. Injection of Cas9 and sgRNA into the zygote marks cells with genetic scars at an early developmental stage. Scars are analyzed by using the CIGAR code, which describes length and position of insertions and deletions. (**b**). Correlation of scar abundances between two different 24 hpf embryos injected with sgRNA and Cas9 mRNA. Each black dot represents a different unique scar. Axes are linear between zero and ten, and logarithmic for higher abundances. Inset: We found 816 and 1175 unique scars in embryo 1 and embryo 2 respectively, compared to 397 scars present in both embryos. (**c**). Dynamics of scar formation after injection of sgRNA and Cas9 mRNA. Fraction of wildtype reads, averaged over multiple embryos, as a function of time. The fitted dynamical model (see also Materials and Methods) suggests Cas9 activity until (10 ± 2) hr. d. Scarring efficiency after injection of uCas9 mRNA, Cas9 mRNA or protein.

In order to determine how many cell lineages can be distinguished with scartrace, we analyzed the complexity of scar sequences in whole embryos at 24 hours post fertilization (hpf). We found that Cas9 generated hundreds of unique scars when targeting a single site in GFP (Fig. 1b, Supplementary Fig. 1, and Supplementary Table 1), suggesting that analysis of genetic scars constitutes a powerful approach for whole-organism lineage analysis. Detection of scar sequences could be performed based on genomic DNA or mRNA, without any apparent differences (Supplementary Fig. 2). Scar abundances spanned several orders of magnitude. While many scars were unique to a particular embryo, we also observed that others, in particular the most abundant ones, appeared in multiple embryos and displayed correlated abundances between different samples. This finding indicated that some scar sequences are more likely to be created than others, probably through mechanisms like microhomology-mediated repair^17^ (Supplementary Fig. 3). Consequently, the scars with the highest intrinsic probabilities will be created multiple times in different embryos.

Scarring continued until around 10 hpf, a stage at which zebrafish already have thousands of cells (Fig. 1c). We used this dynamical data and a mathematical model of scar generation dynamics to approximate the rate at which double strand breaks are induced and to determine the probability α_i_ that a particular scar *i* is generated (Supplementary Fig. 4, see also Materials and Methods). Embryos consistently exhibited a higher percentage of scarred GFP when injecting Cas9 protein compared to Cas9 mRNA, suggesting that protein may act earlier than mRNA (Fig. 1d), leading to larger clone sizes. The dynamics of scar formation can potentially be adjusted further by injecting variants of Cas9. To illustrate this, we constructed an unstable variant of Cas9 (uCas9). Embryos injected with uCas9 mRNA had lower wildtype levels than embryos injected with Cas9 mRNA (Fig. 1d). Thus, our simple injection-based approach for Cas9 induction allowed us to label cells in an important developmental period during which the germ layers are formed and precursor cells for most organs are specified^18^.

We next aimed to use scartrace to determine the clonality of specific cell populations. We chose zebrafish germ cells for this proof of concept experiment, as previous studies have established that the entire germ cell pool is derived from 4 primordial germ cells specified at around the 32 cell stage. These 4 founder germ cells start to proliferate at around the 4,000 cell stage (∼5 hpf) and their total number increases to 25-50 by the end of the first day of development^19^. We hence raised selected histone-GFP zebrafish that were injected with sgRNA and Cas9 protein or mRNA to adulthood. We then bred a heterozygous female with a wildtype male (or vice versa) and sequenced scars in the individual resulting embryos (Fig. 2a). This approach allowed us to sequence the clonal complexity of the maternal or paternal germ cell pool on a single cell level. As expected for a transgene that is integrated on a single chromosome, we found a clear bimodal distribution of GFP expression, with approximately half of the embryos expressing GFP (Supplementary Fig. 5). However, we detected multiple scars per embryo in the F1 generation, indicating that there are multiple integrations of the transgene in the genome. For example, we found embryos derived from a protein-injected parent that have two scars in a 25%-75% ratio, suggesting four integrations. By a similar analysis, we found embryos derived from mRNA-injected parents that suggest eight integrations (Fig. 2b). An analysis of GFP integrations available for scarring over 113 embryos derived from a protein-injected parent, and 397 embryos derived from mRNA-injected parents showed that the average number of GFP integrations available for scarring is (2.9 ± 0.2) for protein injections. For mRNA injections, the average number of GFP integrations available for scarring is (9.4 ± 0.6) (see Materials and Methods). The differences between embryos are likely to be caused by excision of tandem GFP-integrations that result from multiple double-stranded breaks being created at the same time.

**Figure 2.**
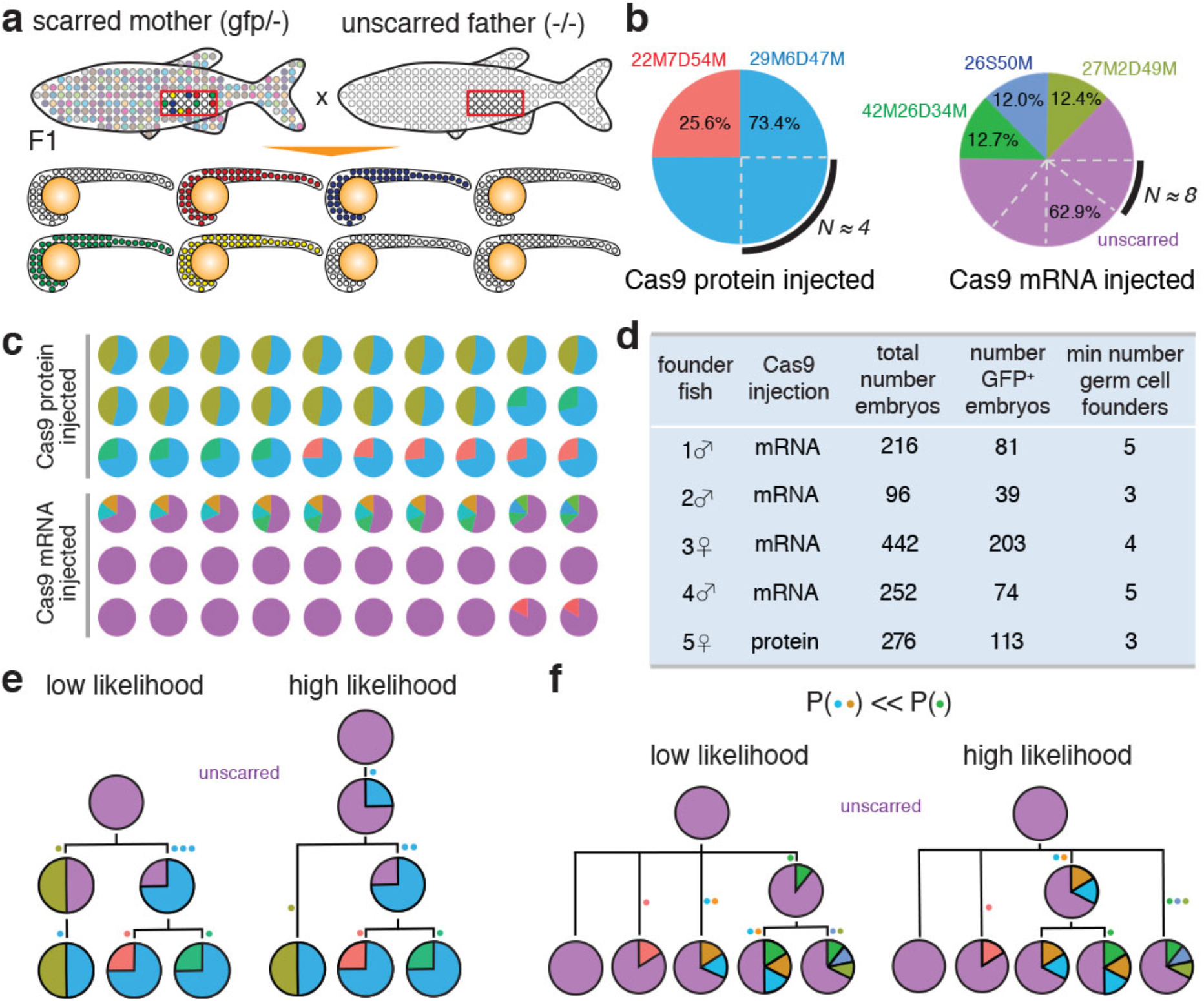
Clonality of the germ cell pool. (**a**). Sketch of the experimental protocol. An adult female with scars induced by injection of sgRNA and Cas9 protein or mRNA was crossed with a wildtype male (or vice versa). Since we used heterozygous fish, only half of the germ cells (red box) carried scars. Cells with different scar profiles are indicated by circles filled with different colors. To analyze the clonality of the scarred parent’s germ cell pool, we sequenced the scars of the F1 generation. (**b**). Representative examples of scar frequencies for two individual embryos derived from a protein and an mRNA-injected parent. (**c**). Representative subset of embryos from the same parents, chosen to show all clones at approximately the ratio in which they are found in the data. (**d**). Overview of the number of embryos and detected clonality of the germ cell pool for five different scarred founder fish. (**e**). Two examples of lineage trees for the F1 clones derived from a protein-injected parent. The tree with the maximum likelihood is shown on the right. The likelihood of the left tree is about 6 fold lower with respect to the maximum likelihood. (**f**). Two examples of lineage trees for the F1 clones derived from a mRNA-injected parent. The tree with the maximum likelihood is shown on the right. The likelihood of the left tree is more than 10^7^ fold lower with respect to the maximum likelihood.

When crossing a founder fish injected with Cas9 protein, we observed the same three dominant scar profiles (Fig. 2c), which suggests that there were at least three independent clones of germ cells. For Cas9 mRNA injected founder fish, we found slightly higher numbers of germ cell clones (Fig. 2c,d), consistent with the notion that Cas9 mRNA acts later than protein. We next used the clones shown in Figure 2c to construct lineage trees of the germ line of a Cas9 protein-injected fish (Fig. 2e) and a Cas9 mRNA-injected fish (Fig. 2f). We used the scar probabilities we previously determined to compare the likelihoods of lineage trees, and selected the tree that is most likely to be correct. We show two possible lineage trees for a protein and mRNA injected founder fish in Figure 2e and 2f respectively. For example, the right lineage tree of the mRNA injected fish (Fig. 2f) is much more likely than the left tree, because the green scar (27M2D49M, α ≈ 0.034) is much easier to create than the orange scar, which is a complex scar consisting of two adjacent deletions (25M3D11M4D40M, α < 10^-5^). Therefore creating an orange scar twice in the same embryo is very unlikely, in contrast to creating multiple green scars.

In summary, these experiments validated scartrace as a reliable and reproducible approach for quantitative whole-organism clonal analysis and showed its applicability for quantitative lineage tracing on select cell populations. The high diversity of scars, combined with variable numbers of integrations available for scarring, created a very high complexity of scar profiles for clonal analysis. Excision of concatemerized target sites has been observed before for Cas9 protein injections^13^. This poses a challenge for lineage tracing, as it leads to the erasure of scars that were previously introduced. For the remaining experiments in Figure 3 we therefore focused on Cas9 mRNA injection experiments, in which target excision is much less likely.

**Figure 3.**
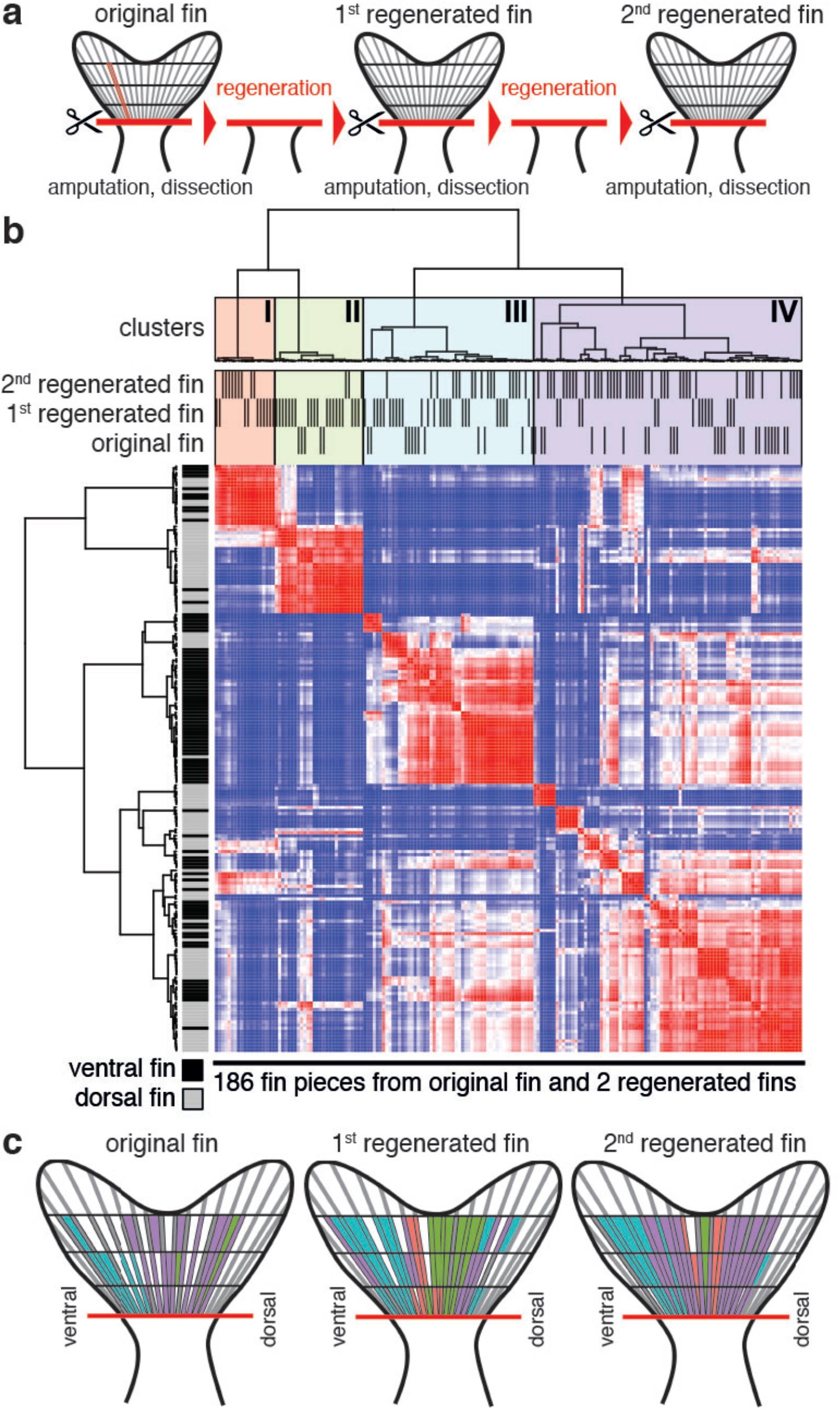
Clonality of the regenerating caudal fin. (**a**). Sketch of the experimental protocol. We amputated the caudal fin of an adult zebrafish that was injected with sgRNA and Cas9 mRNA at the 1-cell stage. The fin was dissected into proximal, central, and distal pieces of rays and interrays, and the scar profile of the individual pieces was analyzed by sequencing. This experiment was repeated twice after regeneration of the fin. (**b**). Clustering of fin pieces by similarity of scar profiles, normalized by global scar probabilities (see also Materials and Methods). Clusters were determined for a combined dataset including all data from the original and regenerated fins (186 pieces in total). The panel below the four clusters indicates from which fin (original, 1^st^ or 2^nd^ regeneration) the individual pieces were isolated. (**c**). Spatial profile of scar clusters. Only pieces for which we detected sufficiently high read counts are shown (see also Materials and Methods). Color code as in Figure 3b.

After this first proof of principle, we decided to apply scartrace to a more complex, yet well-studied biological system. We chose to focus on the zebrafish caudal fin, a structure that consists of about a dozen different cell types. Zebrafish have the remarkable capacity to regenerate fins upon amputation, with most cell types in the regenerated organ derived from cell-type specific precursors^20^. We amputated the caudal fin of an adult fish marked with Cas9-induced scars, and we dissected the fin into pieces consisting of proximal, central, and distal positions of individual rays and interrays (Fig. 3a). This procedure was repeated twice after the fin had fully regenerated. We then subjected the individual pieces of the original and the two regenerated fins to sequencing in order to identify their scar profiles. Each piece consisted of a mixture of different cell types, and we typically detected between 10 and 30 distinct scars per piece (Supplementary Fig. 6). We normalized detected scar abundances by global scar probabilities and performed hierarchical clustering on the data (Fig. 3b, Supplementary Fig. 7, and Materials and Methods). We found that the scar profiles of the different pieces segregated into four clusters. The clusters consisted of a mix of pieces from the original and the regenerated fins, suggesting that the differences in scar profiles within a dissected fin may be bigger than between the original and regenerated fins. The clusters were spatially organized along the dorsoventral and anteroposterior axis (Fig. 3c). Interestingly, the spatial position of the individual clusters remained largely constant between the original and the regenerated fins. Correlation was particularly strong between pieces belonging to the same ray or interray, suggesting that growth patterns predominantly followed the proximal-distal axis (Supplementary Fig. 8). These findings indicate, in agreement with previous reports^20,21^, that formation of the caudal fin proceeds similarly during development and during regeneration. Interestingly, we found that the number of detected scars decreased only mildly in the regenerated fin (Supplementary Fig. 9). This observation suggests that most clones that gave rise to the original fin survived until adulthood and were reactivated upon amputation.

Here we introduced scartrace, a simple approach for systematic lineage analysis of whole organisms and for quantification of the clonality of tissues and organs. A key advantage of our approach is that cell labeling as well as detection of lineage markers can be performed in a high-throughput manner. For instance, previous studies of fin regeneration required manual analysis of hundreds or thousands of fish, in which the progeny of only one or a few cells was traced^20,21^. Our approach allows lineage analysis of many cells in the same organism, facilitating for instance quantitative studies of clonality changes during life and after perturbation. Importantly, scartrace does not require prior knowledge of cell type markers. Scartrace will hence be an ideal method for elucidating the origin of anatomical structures whose developmental lineage is unknown. Injection of Cas9 and sgRNA into the zygote of histone-GFP zebrafish enabled us to label most cells in the organism at important developmental stages such as mid-blastula transition and gastrulation. Scartrace is based on a fish line that is already present in most zebrafish facilities and on easily available reagents, and can hence be adopted immediately by other labs.

Viral barcoding of single cells has been used to track lineages in the hematopoietic system^11,12^, and it should in principle be possible to combine this method with single-cell RNA-seq. One important downside of viral barcoding is that it is difficult to apply the technique efficiently in systems that are less well suited for *ex vivo* infection and transplantation than the hematopoietic system. More fundamentally, viral barcoding is generally limited to a single time point, which precludes reconstruction of full lineage trees. Our strategy is similar to a recently published method using concatemerized Cas9 target sites^13^. Concatemerized target sites have the advantage that multiple scars can be sequenced on the same read. However, a potential downside is that target sites may easily be excised, leading to loss of previously established scars. Cumulative acquisition of scars can be used for reconstructing lineage trees. It is however important to take into account that scars have different intrinsic probabilities, and that the probability for repeated generation of the same scar in a single embryo is sequence-dependent. Using our mathematical model of scar generation dynamics, we quantified these probabilities and incorporated this information into algorithms for lineage reconstruction (Fig. 2e,f) and clustering (Fig. 3b).

We expect that there will be variants of Cas9-mediated lineage tracing in the future, using for instance self-targeting sgRNAs^22,23^ or inducible systems. Combination of scartrace with single-cell RNA-seq will ultimately enable unbiased cell type identification and simultaneous lineage reconstruction of heterogeneous cell populations. Lineage tracing of single cells in the human body will be an important challenge for future research. We anticipate that scartrace and related experimental approaches will allow researchers to refine and validate purely computational lineage tracing methods, which do not require genetic manipulation^24,25^.

## Materials and Methods

### Zebrafish

Embryos of the transgenic zebrafish line Tg((H2Af/va:H2Af/va-GFP)^kca66^ ^16^ were injected at the 1-cell stage with 1 nl Cas9 protein (NEB, final concentration 1590 ng/μl) or mRNA (final concentration 300 ng/μl) or uCas9 (final concentration 300 ng/μl) in combination with an sgRNA targeting GFP (final concentration 25 ng/μl, sequence: GGTGTTCTGCTGGTAGTGGT). The uCas9 was constructed by introducing an 8 amino acid long destruction box used in the zFucci system^26^ into the N-terminus of the pCS2-nCas9n vector. Cas9 mRNA and uCas9 mRNA were in vitro transcribed from linearized pCS2-nCas9n vector (Addgene plasmid # 47929)^14^ using the mMESSAGE mMACHINE SP6 Transcription Kit (Thermo Scientific). The sgRNA was in vitro transcribed from a template using the MEGAscript^®^ T7 Transcription Kit (Thermo Scientific). The sgRNA template was synthesized with T4 DNA polymerase (New England Biolabs) by partially annealing two single stranded DNA oligonucleotides containing the T7 promotor and the GFP binding sequence, and the tracrRNA sequence, respectively^18^.

For caudal fin experiments we manually dissected the amputated fin into small pieces corresponding to proximal, central, and distal parts of rays and interrays, as shown in Figure 3a. Amputation and dissection were repeated after the caudal fin had fully regenerated. The genomic DNA from these pieces was then used to determine scar profiles by sequencing. All studies involving vertebrate animals were performed with institutional approval of the Hubrecht Institute, and were reviewed by the dierexperimentencommissie (DEC) of the KNAW.

### Scartrace protocol

The protocol was performed using either reverse transcribed RNA or genomic DNA as starting material. RNA was extracted from homogenized zebrafish samples with TRIzol reagent (Ambion) according to the manufacturer’s instructions. For DNA extraction we used either TRIzol extraction or SEL buffer + Proteinase K (50 mM KCl, 2.5 mM MgCl2, 10 mM Tris pH 8.3, 0.045% Igepal, 0.045% Tween-20, 0.05% Gelatine and 0.1 mg/ml Proteinase K). For extraction with SEL buffer + Proteinase K, samples were incubated for 1 h at 60 °C and 15 min at 95 °C. Scar detection for the experiments shown in Figure 1b was done on mRNA, all other experiments shown here were performed on genomic DNA. To obtain unbiased measurements of scarring dynamics and efficiency, embryos were collected regardless of GFP expression for the experiments in Figure 1. For the experiments in Figures 2 and 3, we excluded fish that continued to express high levels of GFP, which is a hallmark for unsuccessful injections. No randomization of animals or blinding of group allocations was performed.

GFP sequences were amplified by PCR with primers complementary to GFP, including an 8 bp barcode sequence^27^ and adapter sequences for Illumina sequencing. Samples were then pooled and subjected to magnetic bead cleanup (AMPure XP beads – Beckman Coulter). Finally, sequenceable libraries were generated by a second round of PCR with indexed primers from Illumina’s TruSeq Small RNA Sample Prep Kit. We confirmed successful library preparation by Bioanalyzer (DNA HS kit, Agilent). Samples were sequenced on Illumina NextSeq 500 2×75bp.

### Determination of scar abundance

Sequencing data were mapped to GFP using bwa mem 0.7.10. We only considered reads for which the left mate was mapped in the forward and the right mate in the reverse direction, for which the left mate contained a correct barcode, and for which the right mate started with the primer sequence and had a length of 76 nucleotides. The PCR primer locations were chosen such that the scar can be found in the right mate. To identify different scars, we first classified right mate reads using the CIGAR string, which describes length and position of insertions and deletions in the alignment, combined with the 3’ location of the right mate. Within each CIGAR string we performed a subclassification based on which sequences it contained. We retained those CIGAR strings for which we saw at least 20 reads in a library, and those sequences that made up at least 5% of all reads in its CIGAR-class.

### Dynamical model of scar generation

We assumed that Cas9 is active until time *t*\* and that during this period double-strand breaks (DSB) can be independently generated in unscarred integrations at a rate γ. Accordingly, the average fraction *f_WT_*(*t*) of wild-type integrations at time t is:

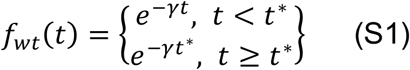

This equation can be fitted to the average fraction of wildtype GFP reads as measured in dynamic experiments performed with mRNA injected fish (Fig. 1c). We obtained *γ* = 0.055 ± 0.003 h^-1^ and *t*\* = 10 ± 2h.

We assumed that scars are formed immediately after Cas9-induced generation of DSBs. We denote *α_i_* the probability of scar *i* to be generated after DSB formation. The average fraction of scar *i* at time *t* is given by:

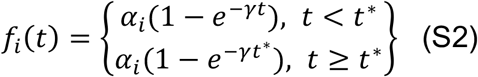

This expression can be fitted for each scar using the dynamic data measured in Figures 1c to obtain the probabilities *α_i_* for each scar. The resulting fits for two different scars are shown in Supplementary Figure 4. Finally, we set the probability of any scar we do not observe in the dynamic experiments to 10^-5^. This corresponds to the average over the different values of *α_i_* obtained for scars that are only observed once in dynamic experiments.

### Microhomology analysis

To determine how many of the scars we observe could be created through microhomology-mediated repair, we simulated this repair mechanism *in silico.* We divided the DNA around the Cas9 target site into two DNA stretches corresponding to a double-stranded break three bases upstream of the PAM sequence, the most frequent cut site. We then removed bases from the 5’ and 3’-stretch, and determined if the loose ends had a microhomology. By doing this for every combination of bases removed from the 5’ and 3’-ends, we created a list of sequences created through microhomology-mediated repair.

We next ordered all observed scars according to their probabilities, and determined which scar was created through microhomology-mediated repair. The cumulative plot shown in Supplementary Figure 3 shows that the most probable scars are enriched in microhomology-mediated repair.

### F1 embryo data analysis

After determining scar abundances as described above, we selected samples with a sufficiently high read count: we first calculated the average number of reads for samples in a given library, and then filtered out all samples that did not have at least 10% of this average. After this, we singled out scars that made up at least 5% of all reads of at least one embryo. The abundances of all other scars were summed up and displayed as ‘other’. We keep embryos that have a scar makeup that is reproduced by at least one other embryo.

To determine the number of GFP integrations available per F1 embryo, we first determined for every embryo the scar with the lowest ratio higher than 5% of all reads in that embryo. The inverse of this ratio is the number of GFP integrations that were available for scarring in the germ cell that gave rise to this embryo. For added precision, we selected per embryo all scars that had ratios similar to the minimum ratio (we defined a similar ratio as a ratio that implied at most one integration more than the minimum ratio did), and took the average over the inverses. We finally calculated the averages over the numbers of GFP integrations for Cas9 protein-injected parents and for Cas9 mRNA-injected parents.

### Lineage tree reconstruction

We constructed F1 lineage trees in a bottom-up fashion, grouping clones together that shared one or more scars, and replacing the clade with an internal state that contains only the shared scars. This yields the trees shown on the right in Figure 2e and 2f. For the trees shown on the left we slightly deviated from this approach to show likelihood ratios between two similar trees.

### Fin data analysis

We selected fin pieces with sufficiently high read count in the same way as we did for the F1 samples. To account for differential scar probabilities, we calculated the ratio between the observed scar probabilities for each fin piece and the global scar probabilities (Supplementary Fig. 4) for each scar. For scars in the fin data that were detected with a probability of less than 10^-5^, the normalization factor was set to 10^-5^. This strategy was successful in removing scar abundance correlations between different embryos (Supplementary Fig. 7). We calculated the pairwise Pearson correlations after removing non-scarred GFP. The hierarchical clustering in Figure 3b was done on pairwise correlations between the pieces, using Ward’s method for agglomeration to find clusters with minimal within-cluster variance. We cut the resulting dendrogram into four clusters that were determined to be stable (with Jaccard similarities ranging from 0.72 to 0.88) even when clustering a subset of fin pieces. We used the function ‘clusterboot’ in the R package ‘fpc’ for this stability assessment.

### Confidence intervals

The 95% confidence intervals shown in Figures 1c, 1d, Supplementary Figure 4 and Supplementary Figure 8, and those for the number of GFP integrations available were generated by sampling the populations with replacement, determining the mean of the population, repeating this thousand times and taking the upper boundary of the lowest 2.5%-quantile and the lower boundary of the highest 2.5%-quantile of all means generated. For the number of GFP integrations, we enlarged the confidence interval to be symmetric around the observed mean. Sample size was 100 for Figure 1c, 40 for Figure 1d, and 203 and 113 for mRNA and protein respectively for the number of GFP integrations available.

